# Multilayered specificity of transcription factor binding at cytokine promoters

**DOI:** 10.64898/2026.06.08.730942

**Authors:** Anna Lagani, Ryan Lane, Yunwei Lu, Sakshi Shah, Zhaorong Li, Luis Soto-Ugaldi, Mahir Patel, Cosmin Ciausu, Matias Alejandro Paz, Juan I. Fuxman Bass

## Abstract

Transcription factors (TFs) regulate gene expression through sequence-specific DNA binding, and their genomic occupancy is further influenced by TF expression, activation state, and protein-protein interactions. How these mechanisms determine context-specific gene regulation remains incompletely understood, particularly for tightly controlled immune genes such as cytokines. Here, we use paired yeast one-hybrid (pY1H) assays to systematically examine DNA binding of 236 TFs and 392 TF-pairs across 106 cytokine gene promoters. Of the 1,619 TF-promoter interactions identified, 555 required TF cooperativity and 410 were antagonized by at least one TF partner, suggesting that TF-DNA binding is highly dependent on TF partners. Usage of different partners can drastically alter a TF’s target repertoire and may result in the recruitment of different transcriptional cofactors. Integration with existing data on TF expression and activation further showed that cooperativity and antagonism provide additional, underappreciated layers of DNA-binding specificity.

## INTRODUCTION

Specificity in gene expression underlies cell type-specific and context-dependent processes such as differentiation^1^, development^2,3^, and responses to environmental or pathogenic stimuli^4-6^. This specificity requires tightly regulated binding of transcription factors (TFs) to cis-regulatory DNA elements (CREs) such as promoters and enhancers. Genetic variation affecting CRE sequence or TF function can broaden, restrict, or alter transcriptional patterns, contributing to diverse diseases including immune disorders, developmental abnormalities, and cancer^7,8^. Determining how various layers of regulation contribute to TF specificity is therefore essential to understand how cellular processes are dynamically controlled at the transcriptional level.

Most studies have emphasized either patterns of TF expression^9^ or control of TF activity^10,11^ - through ligand binding or signaling pathways - as key determinants of TF specificity. However, the mechanisms by which activated TFs select specific DNA targets remain incompletely understood. Work from the past two decades has shown that transcriptional regulation of highly dynamic genes often relies on combinatorial binding of multiple TFs to CREs. This may explain why expression of individual TFs and their target genes are often poorly correlated^12^, and why many ubiquitously expressed and active TFs regulate different sets of genes in different cell types and conditions.

While some CREs may be targeted by multiple TFs binding independently, combinatorial binding can also be facilitated by cooperativity between TFs, often mediated by protein-protein interactions (PPIs)^13^. Cooperativity has the potential to greatly improve TF specificity; the requirement for two or more different TF subunits to bind together generates an AND gate which restricts the opportunities for each TF to bind, and TF-pairs often target DNA sequences that are considerably different from the preferred binding motifs of either TF subunit. Cooperative DNA binding between TFs has been extensively studied for some TF complexes, such as NF-κB^14,15^, basic leucine zipper (bZIP) TFs^16,17^, and nuclear hormone receptors (NRs)^18^. However, systematic studies of TF cooperativity in gene regulatory networks have been restricted due to limitations in existing techniques used to study co-binding of DNA by TF-pairs, which either study one TF-pair at a time, identify short, preferred binding sequences for TF-pairs *in vitro*, or rely on the coincidence of known motifs for multiple TFs within accessible DNA footprints^15,18-22^. We hypothesize that cooperativity plays a broader role in achieving the observed level of TF-DNA binding specificity than is currently appreciated.

To address these limitations, we recently developed paired yeast one-hybrid (pY1H) assays to identify interactions between hundreds of single-TFs or TF-pairs and DNA regions of interest^23^. In this method, yeast carrying a DNA region of interest integrated upstream of two reporter genes (*HIS3* and *LacZ*) express either one or two TFs, each of which is fused to the yeast Gal4 activation domain (AD). In the event of TF binding to the DNA region of interest, the AD will activate reporter gene expression, allowing yeast to grow in the absence of histidine and turn blue in the presence of X-gal. Yeast expressing only one TF are compared to yeast expressing two TFs to identify DNA binding events that require cooperativity. In our pilot screen testing 18 promoter sequences, we observed cooperative binding from a wide variety of TF-pairs in our array as well as DNA-binding antagonism, in which one TF sequesters another to prevent its binding to certain DNA regions^23^.

Here, we greatly expand these preliminary studies to investigate how DNA-binding cooperativity and antagonism contribute to TF specificity along with well-studied regulatory mechanisms, including TF expression and activity modulation. As a model for investigating these layers of specificity, we mapped interactions between TFs and the promoters of cytokine genes. While it is known that many of the ∼130 human cytokines are expressed only in specific cell types and/or in response to cellular signals, it is not fully understood how this high level of specificity is achieved to prevent aberrant immune activation. By studying the binding of single-TFs and TF-pairs to cytokine gene promoters, we greatly expand the cytokine gene regulatory network (GRN), observe extensive combinatorial targeting of cytokine genes by known and novel cooperative TF-pairs, and reveal a widespread role of TF-TF antagonism in specifying cytokine gene targets of TFs. Combining our findings with information about TF expression and activation, we consider these multiple layers of TF specificity in combination, observing that ∼97% of interactions we identified are subjected to at least one of these layers. Importantly, TFs whose interactions with cytokine promoters are regulated by more layers of specificity are more often associated with immune-related phenotypes, suggesting a role for these specificity mechanisms in controlling immune gene expression.

## RESULTS

### Unbiased expansion of the cytokine GRN

Previously, we generated a literature-derived cytokine GRN and the CytReg database, observing a clear bias towards highly studied TFs and cytokines^24,25^. To uncover understudied portions of the GRN, we had used enhanced yeast one-hybrid (eY1H) assays (**Fig. 1a**) to evaluate the binding of 1,086 TFs to promoters of cytokine genes (encompassing 2 kilobases upstream of the transcription start site) (**Supplementary Table 1**), identifying 1,380 protein-DNA interactions (PDIs) between 265 TFs and 108 cytokine promoters^25^. However, eY1H assays test each TF independently, and therefore cannot detect interactions involving TFs that function as heterodimers, including important immune-related TFs such as NF-κB and AP-1. To study the DNA binding of TFs as pairs and reveal interactions that require cooperativity between TF partners, we here used pY1H assays to screen 236 single-TFs (**Supplementary Table 2**) and 392 TF-pair combinations (**Supplementary Table 3**) for interactions with each cytokine promoter in a pairwise manner. This enabled us to observe single-TF, cooperative, and antagonistic DNA binding events (**Fig. 1a**).

**Figure 1.**
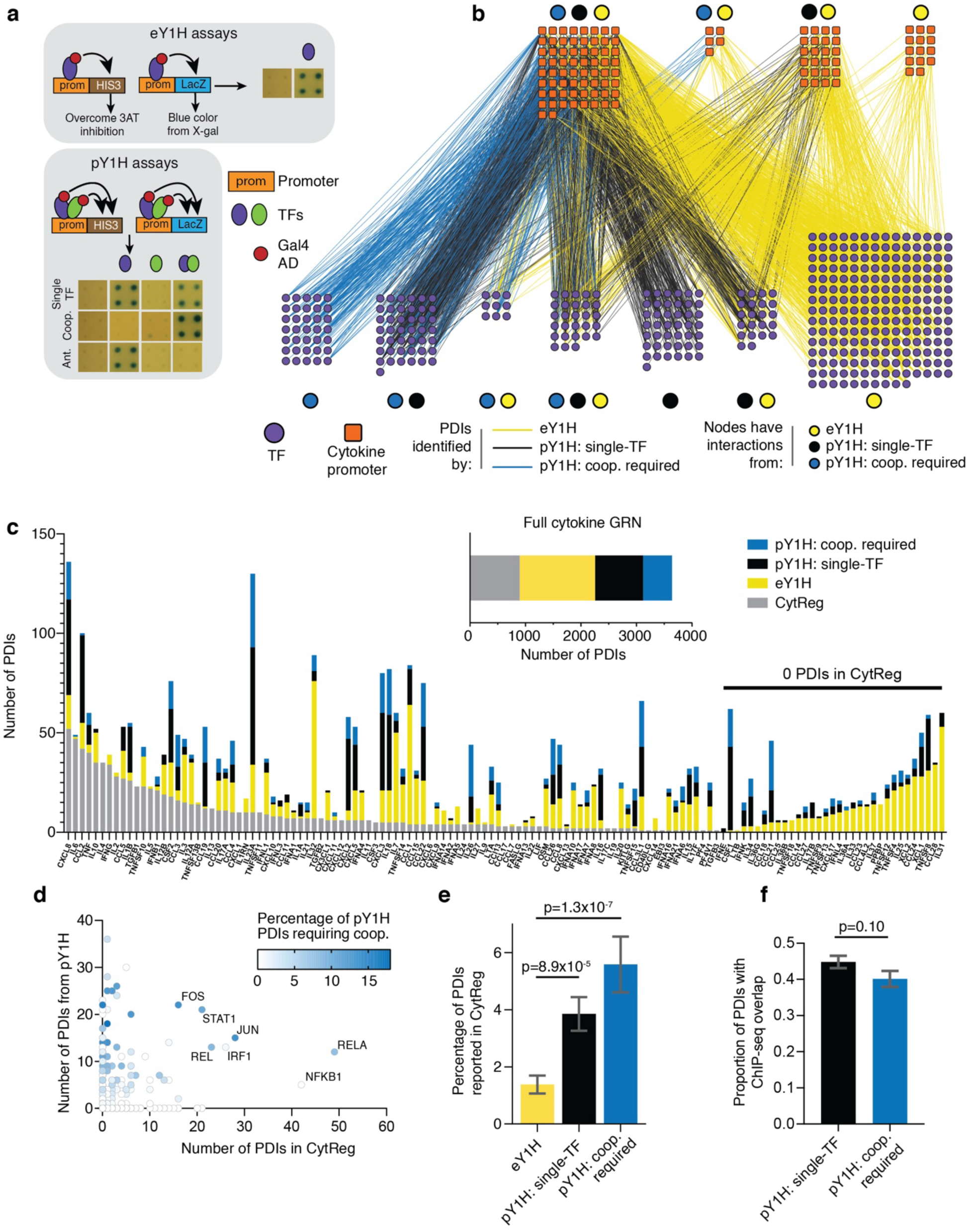
pY1H assays expand the cytokine GRN. **a.** Schematics of eY1H and pY1H assays. eY1H assays detect interactions between single TFs and promoters of interest. pY1H assays can detect single-TF interactions, cooperative binding of TF-pairs, and DNA-binding antagonism between TFs. **b.** The Y1H-derived cytokine GRN, incorporating TF-promoter interactions detected by eY1H and pY1H assays. Purple circle nodes represent TFs and orange square nodes represent cytokine gene promoters. Edges are colored to indicate whether the TF-promoter interaction was identified by eY1H assays (yellow), by pY1H assays as a single-TF interaction (black), or by pY1H assays as an interaction requiring cooperativity with another TF. Nodes are clustered depending on which assays identified interactions involving that node. **c.** Number of PDIs reported for each cytokine promoter in the literature-derived CytReg database, eY1H assays, or pY1H assays. Cytokines are sorted by descending number of interactions reported in CytReg. The inset shows the sum of PDIs for all cytokines from each source. **d.** Number of PDIs in the CytReg and pY1H-derived cytokine GRNs for each TF. Blue shading of each data point corresponds to the percentage of pY1H PDIs involving that TF which required cooperativity with a TF partner. **e.** Percentage of PDIs identified by eY1H (n=1379) and pY1H assays (single-TF n=1064; coop. required n=555) that were previously reported in the CytReg database. Error bars represent the standard error of proportion. Significance by two-tailed proportion comparison test. **f.** Proportion of pY1H-derived PDIs (single-TF n=857; coop. required n=476) with supporting ChIP-seq evidence, defined as the presence of a ChIP-seq peak for binding of the TF at the cytokine promoter in at least one cell type. Error bars represent the standard error of proportion. Significance by two-tailed proportion comparison test.

Using pY1H assays, we identified 1,619 PDIs between 181 individual TFs and 96 cytokine promoters (**Supplementary Table 4**). Importantly, 555 of these PDIs required the coexpression of a TF partner, demonstrating a key role for TF cooperativity in the cytokine GRN. Indeed, in the Y1H-derived network (including eY1H and pY1H interactions) **(Fig. 1b)**, 35 TFs - including immune-related TFs FOSB, RELB, and MYC - only interacted with cytokine promoters via cooperativity and therefore could not have been implicated in cytokine regulation using eY1H assays alone.

Our pY1H-derived interactions expanded the cytokine GRN to a similar degree as the eY1H-derived interactions, and together our Y1H-derived network quadruples the size of the literature-derived CytReg GRN (**Fig. 1b-c, Supplementary Table 4**). We identified novel TF interactions at promoters for cytokines with as many as 57 and as few as zero regulating TFs reported in CytReg (**Fig. 1c**), demonstrating the ability of pY1H assays to expand the cytokine GRN by adding interactions for both highly- and lowly-studied cytokines. Furthermore, pY1H assays identified novel interactions involving well-known immune-related TFs such as NF-κB, AP-1, and STAT family members, while also adding 77 TFs to the network that were not previously associated with cytokine regulation, many of which relied heavily on cooperative binding to target cytokine promoters (**Fig. 1d**). These include members of all major TF families, and 27/77 (35%) have previously been associated with immune-related mammalian diseases annotated in the Mouse Genome Informatics (MGI) database. While expanding the cytokine GRN by adding novel interactions, pY1H-derived PDIs also showed a significantly greater overlap with CytReg than eY1H-derived PDIs (**Fig. 1e**), suggesting that pY1H assays reveal physiologically relevant interactions, and a significantly greater overlap with existing ChIP-seq peaks than expected by chance (**Fig. S1a**), with PDIs that require cooperativity validating at a similar rate to single-TF interactions **(Fig. 1f)**.

Of the 181 individual TFs with pY1H interactions, 24 had been annotated by Lambert *et al.* to bind DNA as “obligate heterodimers,” requiring cooperativity with a TF partner^26^. We therefore expected these TFs to participate exclusively in cooperative binding events rather than single-TF interactions in pY1H assays. However, 15 of these 24 TFs annotated as “obligate heterodimers” exhibited at least one single-TF interaction in the absence of a heterodimeric partner (**Fig. S1b**). This suggests that few TFs are entirely “obligate” heterodimers; rather, many TFs require heterodimerization to bind a subset of DNA targets but can bind elsewhere as monomers or homodimers^21,27^.

### Identification of novel co-binding TF-pairs

The 555 TF-promoter interactions requiring cooperativity derive from 328 cooperative TF1-TF2-promoter binding events between 109 distinct TF-pairs and 73 cytokine promoters (**Fig. 2a, Supplementary Table 5**). These involve TFs across various TF families with no clear enrichment compared to the full TF-pair array (**Fig. S2a**). TF-pairs containing two TFs from the same family (**Fig. 2a**, dark purple nodes) had similar numbers of interactions as pairs containing TFs from different families (**Fig. 2a**, light purple nodes). For example, TF-pairs targeting the largest number of cytokines included GATA1-ZFPM2 (different families), IRF9-STAT1 (different families), FOS-JUN (same family), and ESR2-NCOA1 (different families). This illustrates the prevalence of cooperativity across both same- and different-family pairs. Although most reports have focused on same-family TF heterodimers (e.g. NRs, bZIPs, and basic helix-loop-helix (bHLH), our results are consistent with *in vitro* motif studies using SELEX-seq to test a large, unbiased repertoire of TF-pairs^21^.

**Figure 2.**
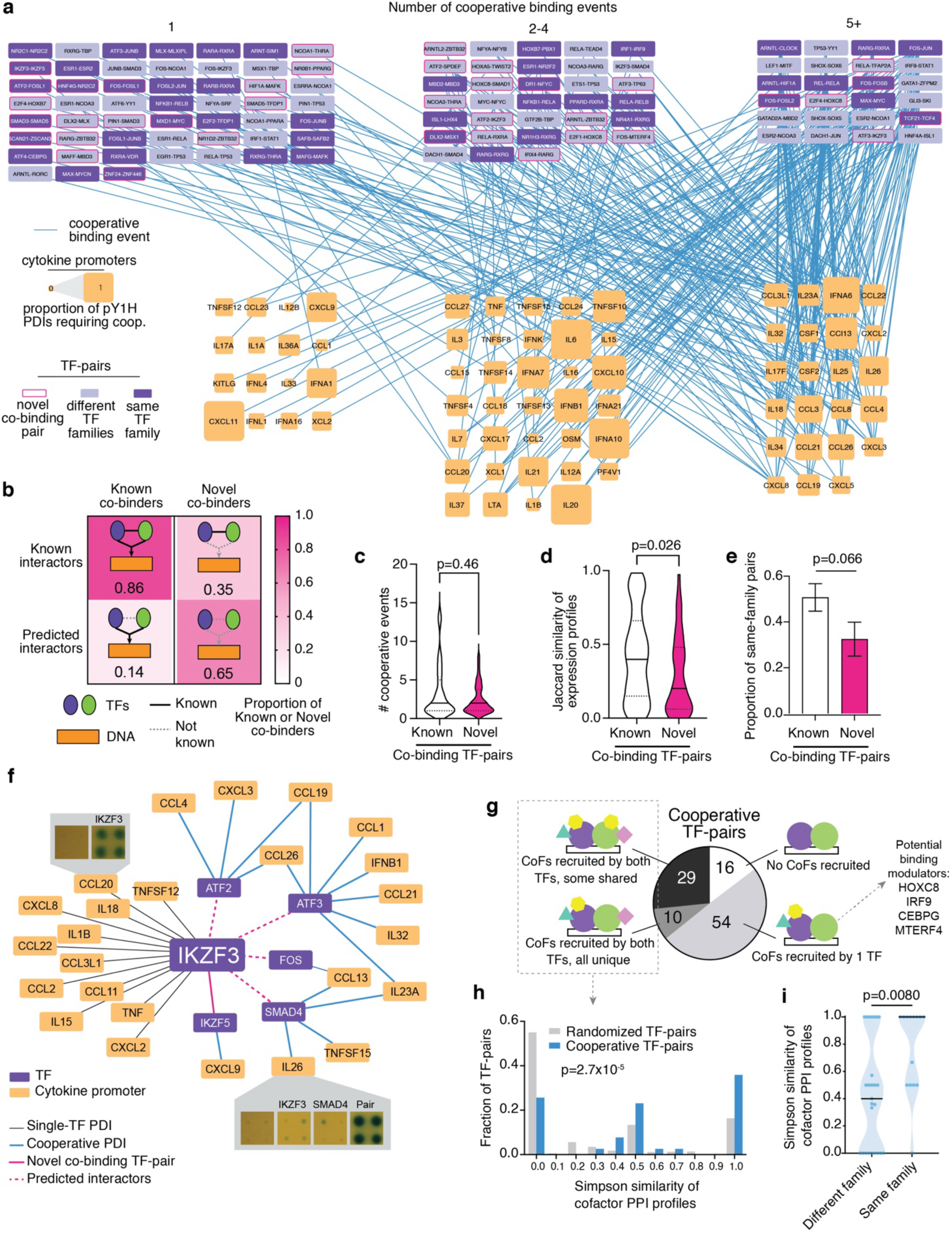
pY1H assays identify cooperative DNA binding of known and novel co-binding TF-pairs. **a.** Network of cooperative binding events between TF-pairs (purple nodes) and cytokine promoters (orange nodes) identified by pY1H assays. The size of the cytokine promoter node corresponds to the proportion of pY1H-derived PDIs at that promoter which required cooperativity between two TFs. TF-pair nodes outlined in magenta represent TF-pairs that have not been previously shown to co-bind DNA (“novel co-binding pairs”). Light purple nodes indicate TF-pairs in which the TFs are members of different TF families, and dark purple nodes indicate TF-pairs in which the TFs are members of the same TF family. Nodes are grouped by the number of cooperative binding events in which they were involved in pY1H assays (1, 2-4, 5 or more). **b.** Proportion of known (n=69) and novel (n=40) co-binding TF-pairs that were either known or predicted to interact by PPIs. **c.** Number of cooperative binding events observed by pY1H assays involving known (n=69) and novel (n=40) co-binding TF-pairs. Significance by Mann Whitney test. **d.** Jaccard similarity of expression profiles for TFs in known (n=67) and novel (n=40) co-binding TF-pairs. Expression levels of each TF across 187 tissue/cell types were obtained from the Tabula Sapiens scRNA-seq atlas. A TF was considered to be expressed in a tissue/cell type if the counts-per-million (CPM) value was at least 10% of the maximum CPM value for that TF across all tissue/cell types. Significance by Mann Whitney test. **e.** Proportion of known (n=69) and novel (n=40) TF-pairs containing TFs from the same TF family. Error bars represent the standard error of proportion. Significance by two-tailed proportion comparison test. **f.** Network of pY1H-derived interactions involving IKZF3. Purple nodes represent TFs and orange nodes represent cytokine promoters. Black edges represent single-TF binding of IKZF3 to cytokine promoters. Blue edges represent cooperative binding of TF-pairs involving IKZF3 to cytokine promoters. Magenta edges connect IKZF3 to cooperative TF partners, all of which form novel co-binding TF-pairs. Example yeast plate images are shown for the single-TF interaction of IKZF3 at the *CCL20* promoter and the cooperative binding event between the IKZF3-SMAD4 pair and the *IL26* promoter. **g.** Number of cooperative TF-pairs in which zero, one, or two TFs are known to recruit transcriptional cofactors. For pairs in which only one TF is known to recruit cofactors, the other TF is considered to potentially function as a DNA binding modulator, rather than a transcriptional effector, if it does not have known transcriptional activating or repressing function. **h.** Distribution of Simpson similarity values for cofactor PPI profiles between TFs in cooperative TF-pairs (n=39) or a set of randomized TF-pairs (n=3900). Significance by Mann Whitney test. **i.** Simpson similarity values for cofactor PPI profiles between TFs in different-family (n=25) or same-family (n=14) cooperative TF-pairs. Significance by Mann Whitney test.

The pY1H array includes: 1) TF-pairs in which the two TFs have been previously reported to interact by PPIs (known interactors), and 2) additional pairs selected based on homology with known interacting pairs (predicted interactors). However, of the 109 TF-pairs that cooperatively bound at least one cytokine promoter, only 69 (63%) have been previously shown to co-bind DNA cooperatively or as heterodimers, while the rest are likely “novel co-binding pairs” (**Fig. 2a**, nodes outlined in pink). Most novel co-binding pairs correspond to predicted, rather than known interactors (**Fig. 2b**); these pairs may interact by previously-unreported PPIs, may form PPIs only once bound to a DNA scaffold, or may cooperatively bind DNA without extensive PPIs between them, as has been reported in previous studies^21^. Importantly, known and novel co-binding TF-pairs participated in similar numbers of cooperative binding events (**Fig. 2c**), suggesting that these newly identified pairs contribute comparably to cytokine promoter occupancy.

We assessed whether novel co-binding pairs may have been understudied due to limited coexpression of the two TFs across tissues and cell types. Using the Tabula Sapiens single-cell (sc)RNA-seq atlas^28^, we mapped the expression of each TF across molecularly defined cell types – most of which were immune cells - found in 23 different tissues **(Supplementary Table 6)**. For each TF-pair, we then determined the extent of expression overlap for the two TFs by calculating the Jaccard similarity of their tissue/cell type expression profiles. We observed that novel TF-pairs had a similar degree of expression overlap as known pairs, suggesting that novel pairs have a similar opportunity to co-bind DNA *in vivo* (**Fig. 2d**). TFs in known and novel co-binding pairs are distributed similarly across TF families (**Fig. S2b).** However, compared to known TF-pairs, novel pairs were less likely to involve members of the same family (**Fig. 2e**), consistent with the general bias for studying same-family TF interactions. This further indicates that cooperative binding extends beyond known TF heterodimers, necessitating large-scale studies to determine the prevalence of these binding events among TF-pairs not previously known to interact with each other or co-bind DNA.

The dependency of a TF on cooperative TF partners to bind DNA often varies depending on the target sequence. For example, IKZF3 (Aiolos) targeted 12 cytokine promoters on its own and 13 cytokine promoters by cooperating with five different TF partners (ATF2, ATF3, FOS, SMAD4, and IKZF5) (**Fig. 2f, S2c**). IKZF3 has only been reported to interact with one of these partners (IKZF5), and has not been shown to co-bind DNA with any of them. IKZF3 is known to bind DNA as a heterodimer with IKZF1 (Ikaros) and modulate B lymphocyte development^29-31^. Given the broad expression of IKZF3 in other immune cells, our results suggest that IKZF3 may also partner with other TFs to regulate cytokine expression beyond the B cell compartment.

### Cooperative TF binding may amplify and diversify cofactor recruitment

Cooperative binding of TFs to DNA may not only contribute to improved specificity in target selection, but may also diversify cofactor recruitment patterns and subsequent regulatory outcomes. To explore this possibility, we identified the cofactors that interact with each TF (including interactions with ≥3 pieces of PPI evidence in the BioGRID database^32^) **(Supplementary Table 7)**. For 54/109 of the cooperative TF-pairs, only one of the two TFs has known cofactor interactions **(Fig. 2g)**. This suggests that, for these pairs, one TF may function as a transcriptional effector by recruiting cofactors to DNA, while the other TF acts predominantly as a modulator of DNA binding. Indeed, four of the cooperative TFs that are not known to recruit cofactors (HOXC8, IRF8, CEBPG, and MTERF4) do not have annotated transcriptional effector domains (i.e. activator, repressor, or bifunctional domains), and may therefore function as DNA binding modulators rather than transcriptional effectors.

For 39/109 cooperative TF-pairs, both TFs have reported interactions with cofactors **(Fig. 2g)**. For most of these pairs (29/39), the two TFs share at least one cofactor. We compared the cofactor PPI profiles for the two TFs within a pair by calculating a Simpson similarity index. Overall, cooperative TF-pairs had more similar cofactor PPI profiles than a randomized set of TF-pairs **(Fig. 2h, S2d)**, indicating that cooperative TFs recruit the same cofactors more often than expected. This suggests that cooperative DNA binding may allow for synergistic cofactor recruitment by increasing affinity or enabling the recruitment of multiple cofactor molecules, amplifying the transcriptional signal.

We also observed that cooperative pairs of TFs from the same family have more similar cofactor PPI profiles than TF-pairs from different families (**Fig. 2i**). In this way, different-family TF-pairs may increase the variety of cofactors recruited to a given target locus. Altogether, we observe that cooperative DNA binding has the potential to both amplify the effect of a given cofactor and coordinate the recruitment of multiple distinct cofactors to target DNA regions.

### Promiscuous TFs use distinct partners to diversify promoter targeting and regulatory function

Of the 117 TFs participating in cooperative binding events, 80 were tested with multiple partners Of these, 50 (62.5%) cooperated with two or more TF partners and were considered “promiscuous” cooperative TFs. This is likely an underestimate, as we only tested a subset of possible TF-pairs and some TFs may exhibit promiscuous cooperative binding at other promoter regions not tested here.

We hypothesized that a promiscuous TF may use different cooperative partners to target different promoters, to target promoters in different cell types, or to elicit different effects on transcriptional outcomes **(Fig. 3a)**. For example, IKZF3 cooperated with different TF partners to bind distinct sets of promoter targets (**Fig. 2f**). More generally, we identified a significant correlation between the number of cooperative partners and the number of cytokine promoters targeted by each TF, suggesting that high promiscuity may broaden the repertoire of promoter targets (**Fig. 3b**). We next assessed whether promiscuous TFs interact with each cytokine promoter using one or multiple cooperative partners (**Fig. 3c**). Promiscuous TFs generally interacted with most of their cytokine promoter targets using only one cooperative partner. This shows that promiscuous TFs generally use distinct partners to target different DNA sequences, rather than redundantly targeting each promoter using multiple partners. For example, MYC bound seven promoters by cooperating only with its partner MAX, three promoters by cooperating only with NFYC, and one promoter by cooperating with either MAX or MXD1 (**Fig. 3d**). Importantly, the well-known cooperative TF FOS was the only promiscuous TF to show a preference for redundant binding using multiple TF partners. However, these partners tend to have different effector functions, suggesting FOS-mediated cooperativity provides regulatory rather than DNA binding diversification.

**Figure 3.**
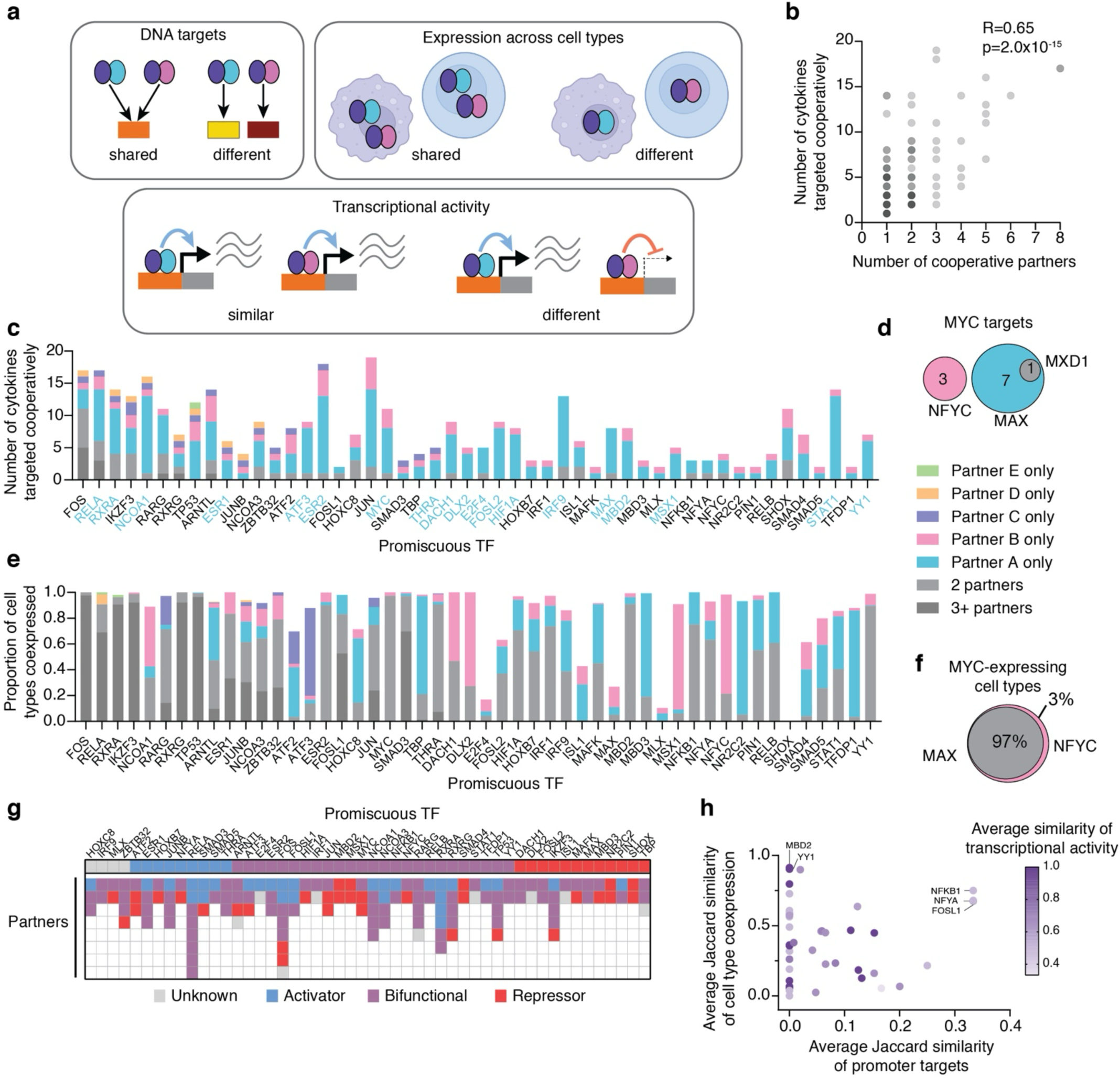
Promiscuous TFs cooperate with multiple TF-partners to expand their DNA target repertoire. **a.** Schematic of potential outcomes for promiscuously cooperative TFs. **b.** Number of cooperative partners and the number of cytokine promoters targeted cooperatively for each TF with ≥1 cooperative binding event observed by pY1H assays (n=117). Significance by correlation test. **c.** Number of cytokine promoters targeted cooperatively by promiscuous TFs. Different colored bars differentiate between promoters targeted using only one TF partner (cyan, pink, periwinkle, peach, and green), two partners (light grey), or three or more partners (dark grey). Promiscuous TFs labeled in cyan have a primary cooperative TF partner. **d.** Number of cytokine promoters targeted by promiscuous TF MYC using TF partners NFYC, MAX, and MXD1. **e.** Proportion of cell types in which each promiscuous TF is coexpressed with its cooperative TF partners. Different colored bars differentiate between cell types in which the promiscuous TF is coexpressed with only one TF partner (cyan, pink, periwinkle, peach, and green), two partners (light grey), or three or more partners (dark grey). **f.** Proportion of cell types in which promiscuous TF MYC is coexpressed with TF partners, NFYC, MAX, and MXD1. **g.** Annotated transcriptional activity of promiscuous TFs and their partners. “Unknown” TFs contain no annotated activation, repression, or bifunctional domains. “Activator” TFs contain only activation domains. “Bifunctional” TFs contain both activation and repression domains or contain at least one bifunctional domain. “Repressor” TFs contain only repression domains. **h.** Summary of the similarity in promoter target repertoire, cell type coexpression patterns, and transcriptional activity between TF partners of each promiscuous TF. Each data point represents one promiscuous TF.

In addition, for about half of promiscuous TFs, we observed a skew where one major “primary” partner was responsible for most of its cooperative interactions (**Fig. 3c**, TFs labeled in blue). Interestingly, the primary cooperative partner of a TF was not necessarily a known DNA co-binder, as known and novel co-binding TF-pairs were equally distributed between primary and secondary cooperative pairings **(Fig. S3a)**. Specifically, ATF3, DLX2, E2F4, FOSL2, MSX1, and THRA all had primary cooperative TF partners with which they had not been previously shown to co-bind DNA, while many known partners of these TFs facilitated few cooperative interactions **(Fig. S3b)**. Therefore, many novel co-binding pairs identified in this study may play a major role in dictating the DNA targets of promiscuous TFs.

Since we observed that different TF partners enable promiscuous TFs to target distinct sets of promoters, we hypothesized that these partners may be expressed in different cell types, facilitating cell type-specific cytokine expression patterns. To test this, we used the Tabula Sapiens scRNA-seq atlas expression data. Most promiscuous TFs were coexpressed with multiple TF partners in the majority of cell types **(Fig. 3e)**. For example, MYC is commonly coexpressed with both MAX and NFYC, although MAX and NFYC facilitate the cooperative binding of MYC to distinct sets of promoters (**Fig. 3f**). This suggests that usage of multiple TF partners typically expands the DNA target repertoire of a TF rather than facilitating DNA binding in different cell types.

We also hypothesized that promiscuous TFs may cooperate with different partners to switch between activation and repression of target gene expression. We therefore classified promiscuous TFs and their partners according to their transcriptional effector domains (activation, repression, and bifunctional domains)^33,34^. TFs with only activation domains were considered transcriptional activators, those with only repression domains were considered transcriptional repressors, and those with both activation and repression domains or with at least one bifunctional domain were considered bifunctional. TFs with no known effector domains were classified as “unknown.” Of the 50 promiscuous TFs, 38 (76%) cooperated with TF partners from at least two different classes (**Fig. 3g**). While most TFs were classified as bifunctional due to the presence of both activation and repression domains, six TFs - SMAD3, ATF3, FOS, RXRG, TP53, and IKZF3 - cooperated with both a putative activator and a putative repressor, suggesting the potential for dramatically different transcriptional outcomes depending on which partner is selected. Even in cases where a promiscuous TF cooperates with multiple partners from the same class, those partners may exert different effects on transcription due to the recruitment of different cofactors, different propensities for forming transcriptional condensates, or different transcriptional kinetics.

We then investigated the relationship between target selection, expression patterns, and transcriptional activity for the partners of each promiscuous TF. For target selection, we compared the set of promoter interactions facilitated by each TF partner by calculating their Jaccard similarity (1 = identical, 0 = no overlap). Similarly, for expression patterns, we compared the sets of cell types where the promiscuous TF is coexpressed with each partner using the Jaccard similarity. Finally, we compared the transcriptional activity of cooperative partners for a given promiscuous TF (1 = same transcriptional effect, 0 = opposite effects). For promiscuous TFs with more than two cooperative partners, we calculated the similarity scores between each combination of TF-pairs and calculated the mean value. For every promiscuous TF, its partners showed major differences in either DNA target selection, expression profiles, or transcriptional activity (**Fig. 3h**). For example, the partners of NFYA, NFKB1, and FOSL1 showed relatively high target similarity and a high level of expression similarity, but differences in transcriptional activity. Conversely, partners of YY1 and MBD2 had very similar expression profiles and transcriptional activity, but facilitated interactions with completely distinct sets of promoters. Altogether, this suggests that the ability to bind cooperatively with multiple partners often provides added specificity in target gene regulation, rather than conveying redundancy to buffer against perturbations in individual TF partners.

### Widespread antagonism between TFs at cytokine gene promoters

In addition to cooperative binding, pY1H assays also detect antagonism, where one TF prevents the binding of another TF. This is identified when yeast expressing only one TF display binding signal, while yeast expressing both TFs show signal loss. Alternatively, if one TF binds regardless of the expression of a second TF, this indicates independent promoter binding (**Fig. 1a**). Of the interactions tested for antagonism, 410 were antagonized by at least one partner (**Supplementary Table 8**), while 305 were independent of all TF partners (**Fig. 4a, Supplementary Table 9**). We observed 175 PDIs that were antagonized by some TF partners but independent of others, suggesting that these PDIs may be either “allowed” or prevented depending on the repertoire of partners present in a given cell type and condition (**Fig. S3c**). Across the cytokine promoters tested, antagonized PDIs were as common as those requiring cooperativity, and combining these two groups, 76% of PDIs detected were dependent on other TFs. This is likely an underestimate of the prevalence of TF-TF dependency, as TFs were tested with only a subset of all possible partners. This constitutes an important paradigm shift in which DNA-binding specificity is not only achieved by cooperative binding but also largely by antagonism between TFs.

**Figure 4.**
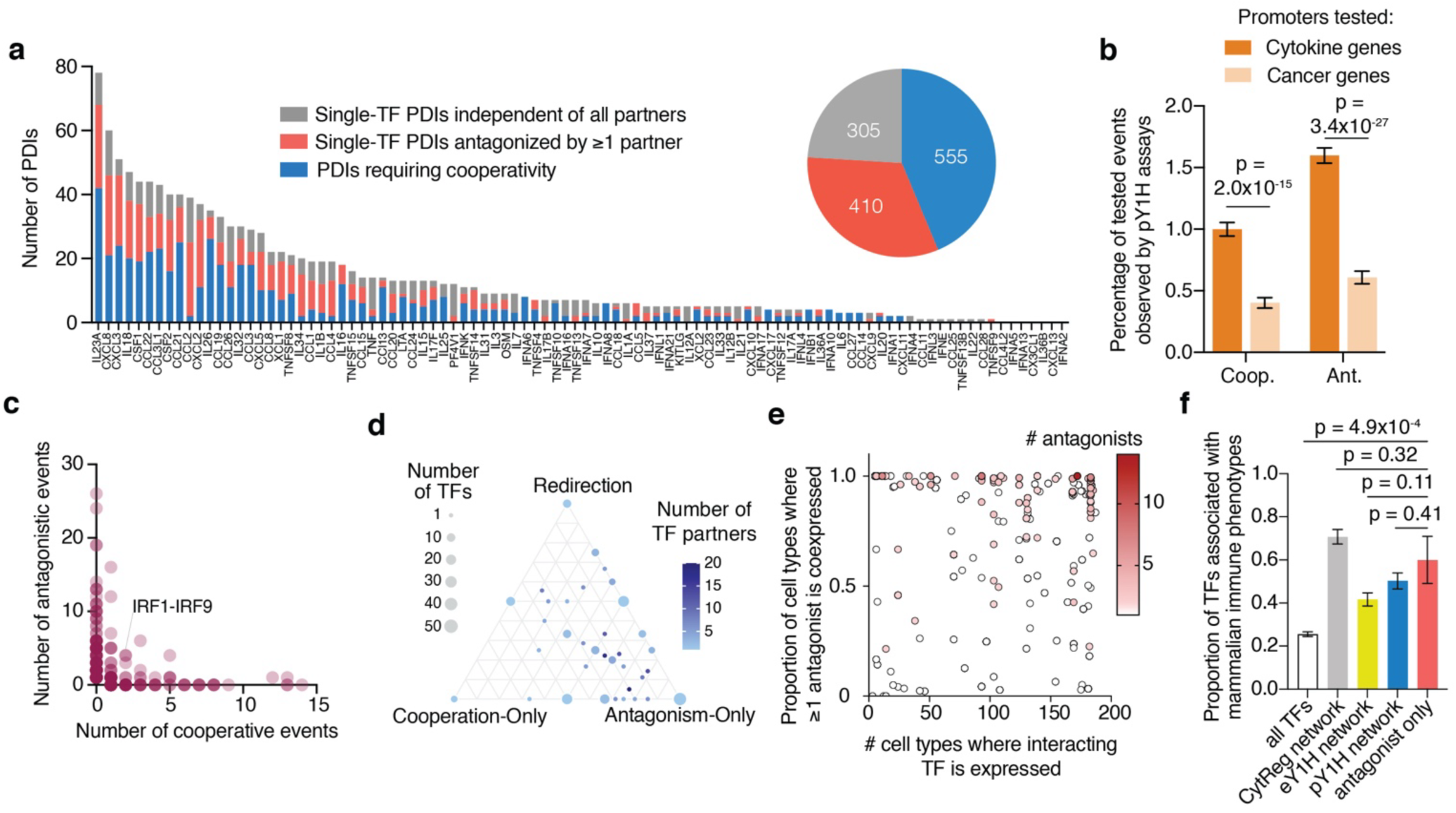
A prominent role for TF-TF antagonism in the cytokine GRN. **a.** The number of PDIs in the pY1H-derived GRN that were independent of all TF partners, antagonized by one or more partners, or required cooperativity for each cytokine promoter (bar graph) and across all promoters (pie chart). **b.** Proportion of all possible cooperative and antagonistic binding events identified by pY1H assays at promoters of cytokine genes (n=32,849 possible cooperative events, n=41,944 possible antagonistic events) and cancer-related genes (n=22,386 possible cooperative events, n=22,386 possible antagonistic events). Error bars represent the standard error of proportion. Significance by two-tailed proportion comparison test. **c.** Number of cooperative binding events and binding antagonism events identified by pY1H assays for each TF-pair. Each data point represents a TF-pair with at least one cooperative or antagonistic event detected by pY1H assays. **d.** Proportion of TF-TF relationships which resulted exclusively in cooperative DNA binding (“Cooperative-Only”), exclusively in antagonism (“Antagonism-Only”), or in both cooperative and antagonistic events depending on the promoter sequence (“Redirection”). Only TFs with 2 or more pY1H interactions are represented. **e.** Number of cell types in which a DNA-binding TF is expressed and the proportion of those cell types in which at least one antagonistic TF is also expressed. Each data point represents a PDI between a TF and a cytokine promoter identified by pY1H assays. Shading of each data point indicates the number of TFs that antagonized the PDI in pY1H assays. **f.** Proportion of TFs associated with mammalian immune phenotypes in the MGI database. Error bars represent the standard error of proportion.

Interestingly, compared to our previous pY1H analyses of promoters from cancer-related genes^35^, cytokine promoters exhibited significantly higher frequencies of both cooperative and antagonistic TF interactions (**Fig. 4b**). This is consistent with the requirement for highly cell type- and condition-specific expression of cytokine genes compared to cancer-related genes, many of which control ubiquitous processes such as cell cycle, transcription, and DNA damage repair.

In the native cellular context, antagonism could have two potential outcomes: either the resulting dimer has no DNA binding ability^36,37^, or it has different DNA sequence specificity^18^ and may bind elsewhere in the genome. In the latter case, antagonism can instead be considered as “redirection” of a TF from one set of DNA targets to another. Indeed, we observed 45 distinct TF-pairs for which one TF antagonized the binding of another to at least one promoter, but both bound cooperatively to at least one different promoter **(Fig. 4c)**, suggesting potential redirection. For example, IRF1 and IRF9 mutually antagonize one another at the promoters of chemokines *CCL21*, *CCL22*, and *CCL3L1*, and IRF1 is further antagonized by IRF9 at the *KITLG* promoter, but the two TFs cooperatively bind the promoters of type-I interferons *IFNA6* and *IFNA7*. This suggests that coexpression of IRF1 and IRF9 promotes their redirection away from promoters of chemokine genes and towards genes involved in the type-I interferon response.

To compare the prevalence of different types of TF-TF relationships, we classified each TF-pair as having an exclusively cooperative relationship (“cooperative-only”), an exclusively antagonistic relationship (“antagonistic-only”), or a relationship that is cooperative at some promoters and antagonistic at others (“redirection”). For each TF, we determined the proportion of its TF-TF relationships that fell into each category. Most TFs participated in multiple types of relationships with different partners, although antagonistic-only pairs were more frequent **(Fig. 4d)**. This suggests that, at cytokine promoters, TFs generally participate in more antagonistic relationships that prevent TF-DNA binding, rather than being redirected to other cytokine promoters.

We hypothesized that DNA-binding antagonism restricts the range of cell types in which a TF can bind its DNA targets, promoting cell type specificity for TF-promoter interactions. To test this, we used the Tabula Sapiens scRNA-seq atlas to determine the set of cell types in which each DNA-binding (antagonized) TF is expressed, representing the full repertoire of contexts where the TF could, in principle, bind DNA. We then identified the proportion of those cell types in which at least one antagonist TF is also expressed for each PDI. Regardless of how broadly a DNA-binding TF is expressed, most PDIs have the potential to be antagonized across the majority of those cell types due to the expression of at least one antagonist **(Fig. 4e)**. As expected, PDIs targeted by multiple antagonistic TFs tended to show higher proportions of cell types with at least one expressed antagonist (**Fig. 4e**, dark red points). Nonetheless, even a single antagonistic partner can substantially constrain the number of cell types in which a TF can engage its target promoters.

Whether a PDI is actually antagonized in a given cell type depends on the relative abundances of the DNA-binding TF and its antagonists. Indeed, we observed that the balance in expression between a TF and its antagonists varied widely across TF-pairs and cell types **(Fig. S3d)**. While some TF-pairs showed a preference for higher expression of either the DNA-binding TF or the antagonist TF across most cell types, others displayed an expression balance that was highly cell type-dependent. This constitutes an additional level of control in determining whether the DNA-binding TF is ultimately available to bind its target, which will also depend on the relative binding affinities of the various components, the repertoire of other TF partners available in each cell type, and the activation state and subcellular localization of each TF.

To determine whether antagonist TFs play a role in immune function, we used the MGI database to identify associations between TFs and either immune-related mammalian phenotypes or human immune diseases. TFs that only acted as antagonists were equally as likely to be associated with immune phenotypes (**Fig. 4f**) and diseases (**Fig. S3e**) as TFs that directly target cytokine promoters **(Supplementary Tables 10-11)**. For example, NR1H2 (also known as LXRB) antagonized RXRG but has not been reported in CytReg, eY1H assays, or pY1H assays to directly target cytokine genes. However, NR1H2 is known to modulate macrophage function, and NR1H2 knockout in mice is associated with increased susceptibility to bacterial infection^38^. Although NR1H2 is known to dimerize with retinoid X receptors like RXRG and modulate the transcription of their targets^39,40^, our results suggest that additional regulation may occur by NR1H2 antagonizing the DNA binding of RXRG, rather than by cooperativity **(Fig. S3f)**. Overall, we propose that TF-TF antagonism plays an important role in regulating the expression of immune-related genes, and that immune disorders may arise from disruptions of antagonist TFs just as they arise from disruptions of DNA-binding TFs.

### The updated cytokine GRN contains both ligand-modulated and cell type-specific TFs

Cooperativity and antagonism between TFs may complement other more well-documented mechanisms for regulating TF function to impart specificity. The transcription of cytokines and other immune response genes has been considered to be predominantly regulated by modulating the activity of ubiquitously expressed TFs. Specifically, the nuclear localization, DNA binding, or transcriptional activity of TFs can be regulated by post-translational modifications (PTMs) or processing, ligand binding, or interactions with other proteins. For example, many immune-response genes are activated by phosphorylation-mediated nuclear translocation of STAT^41,42^ and IRF^43^ TFs, as well as the degradation of I*κ*B to relieve cytoplasmic sequestration of NF-κB dimers^14^.

To determine the contribution of TF activity modulation to cytokine regulation on a broad scale, we annotated the TFs in the cytokine GRN whose activity is controlled by various mechanisms (**Supplementary Table 12**). While 72% of TFs in the literature-derived CytReg GRN are activity-modulated, mainly by PTMs, only 50% of TFs in the full cytokine GRN (CytReg+eY1H+pY1H) are activity-modulated **(Fig. 5a)**. This suggests that TF activity modulation is less predominant than previously assumed, and may reflect a bias towards studying TFs downstream of key immune pathways. Importantly, ∼17% of TFs in the pY1H network are modulated by ligand binding, most of which are in the NR family, suggesting that the expression of many cytokines may be regulated by hormones, metabolites, drugs, and xenobiotic compounds and potentially manipulated using small molecules targeting ligand-binding TFs, as we and others have shown for IL10, CCL2, CXCL8, and IL6^25,44-47^.

**Figure 5.**
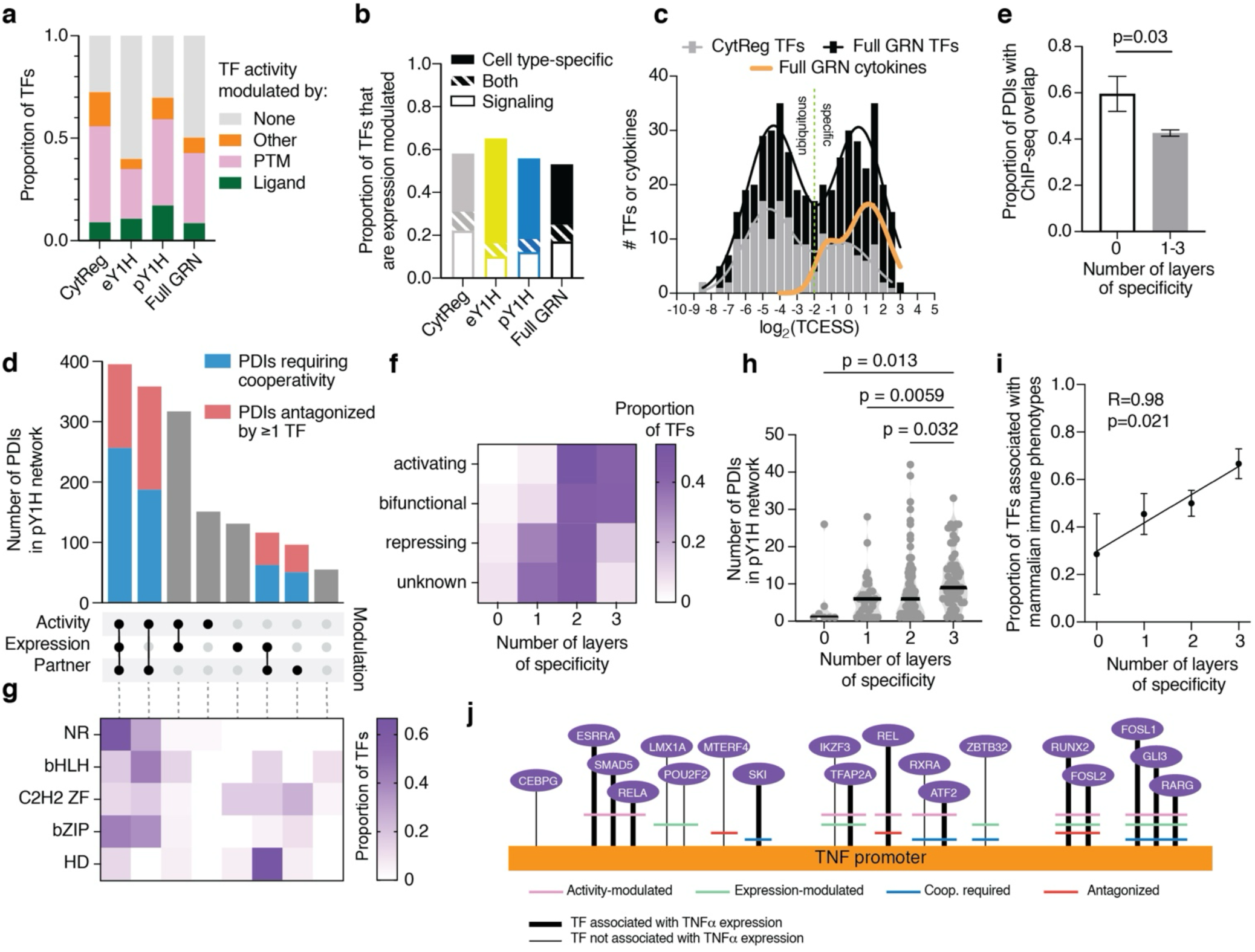
The PDIs in the cytokine GRN are restricted by activity-, expression-, and TF partner-mediation. **a.** Proportion of TFs in the CytReg, eY1H, pY1H, and full cytokine GRNs whose DNA binding activity is modulated by ligand binding, PTMs, or other mechanisms. **b.** Proportion of TFs in the CytReg, eY1H, pY1H, and full cytokine GRNs whose expression is cell type-specific, modulated by signaling pathways, or both. **c.** Histogram showing the frequency of log2(TCESS) values across TFs in the CytReg GRN, TFs in the full cytokine GRN, and cytokine genes. **d.** Number of PDIs in the pY1H network that are modulated at the activity, expression, and/or TF partner levels. **e.** Proportion of pY1H-derived PDIs with 0 (n=42) or 1-3 (n=1291) layers of specificity (activity, expression, and/or partner modulation) with existing ChIP-seq evidence. Error bars represent the standard error of proportion. Significance by two-tailed proportion comparison test. **f.** Proportion of TFs in the pY1H GRN with activating, bifunctional, repressing, or unknown transcriptional activity which are modulated by 0,1, 2, or 3 specificity layers (activity, expression, and/or partner modulation). **g.** Proportion of TFs in the pY1H GRN in the NR, bHLH, C2H2 ZF, bZIP, and HD families which are modulated by different combinations of specificity layers (activity, expression, and/or partner modulation). **h.** Number of pY1H-derived PDIs for TFs modulated by 0 (n=7),1 (n=33), 2 (n=84), or 3 (n=57) layers of specificity. Significance by ANOVA with Kruskal-Wallis multiple comparisons test. **i.** Proportion of TFs in the pY1H GRN modulated by 0 (n=7),1 (n=33), 2 (n=84), or 3 (n=57) specificity layers (activity, expression, and/or partner modulation) which are associated with mammalian immune phenotypes in the MGI database. Proportion by correlation test. **j.** Interactions between TFs and the *TNF* promoter identified by pY1H assays. TFs are arranged from left to right by increasing number of specificity layers modulating each interaction. Thick interaction edges connect TFs that are associated with TNF expression.

In addition to activity-modulated TFs, many GRNs rely on TFs expressed in specific cell types or conditions. To determine the expression specificity of TFs in the cytokine GRN, we first used the Tabula Sapiens scRNA-seq atlas to assign each TF a tissue/cell type expression specificity score (TCESS), where TFs with a TCESS>0.25 are considered cell type-specific^9^ (**Supplementary Table 12**). Second, we used the CytoSig database^48^ to identify TFs whose expression can be modulated by extracellular signaling. TFs meeting either criteria were considered “expression-modulated.” While the CytReg-, eY1H-, and pY1H-derived networks involve similar proportions of expression-modulated TFs, the eY1H network contains a greater proportion of cell type-specific TFs (p=4.10x10^-5^), and the eY1H and pY1H networks both contain a lower proportion of TFs whose expression levels are signal-modulated (p=6.87x10^-5^ and p=0.00499, respectively) **(Fig. 5b)**. Indeed, TFs in CytReg show a bias towards ubiquitously expressed TFs, whereas the full GRN displays a more equal distribution of ubiquitous and cell type-specific TFs, similar to that observed by Ravasi *et al.*^9^ for all human TFs **(Fig. 5c)**. This suggests that the literature bias towards ubiquitous TFs may result from the limited number of studies focusing on tissue or cell type-specific TFs on cytokine regulation, and that expression-modulated TFs may play an critical but underappreciated role in cytokine regulation. More importantly, this may partially explain the high level of tissue/cell type specificity of cytokine genes (**Fig. 5c, green curve**) which was challenging to reconcile with the ubiquitousness of the TFs reported in CytReg. Indeed, of 1,580 interactions in the full GRN involving cell type-specific TFs, 1,458 (92%) connect a TF and target cytokine coexpressed in at least one tissue/cell type, representing possible venues for regulation of cytokines by expression-modulated TFs.

### Multi-layered specificity in TF-DNA binding

Activity modulation, expression modulation, cooperative DNA binding, and antagonism between TFs are all mechanisms that potentially dictate when and where TF-DNA interactions can occur. Considering these mechanisms in combination may help explain how TFs achieve a high level of spatiotemporal DNA-binding specificity despite the prevalence of binding motifs throughout the genome. In particular, the addition of cooperativity and antagonism information may explain how a TF can be expressed and active across multiple cell types, but may target different subsets of genes in those cell types.

To assess the extent to which each PDI in our pY1H network may be limited by each of these specificity mechanisms, we combined literature-derived information about TF activity and expression modulation (**Supplementary Table 13**) with cooperativity and antagonism results from our pY1H assays. For each PDI, we determined whether the TF is activity-modulated (by PTMs, ligand binding, or other mechanisms) or expression-modulated (in certain cell types or stimulatory conditions), and if the PDI is “partner-modulated” (by cooperativity or antagonism). Of the 1619 PDIs in the pY1H network,1564 (∼97%) are potentially restricted by at least one of these three specificity mechanisms (activity, expression, or partner modulation) (**Fig. 5d**), suggesting that most PDIs may only occur under limited conditions. As expected, PDIs with at least one “layer” of specificity were significantly less likely to have been observed in cells by ChIP-seq (**Fig. 5e**), suggesting that these layers effectively limit TF-DNA interactions *in vivo* and therefore reduce the probability of observing these interactions in a given cell type. Interestingly, PDIs subject to activity modulation are typically also subject to expression modulation and/or partner modulation. This may help explain how TFs can be activated by systemic signals (e.g. circulating hormones or cytokines) but only bind DNA targets in specific cell types (i.e., the limited cell types in which the TF is expressed, cooperative partners are also expressed, and/or antagonists are absent).

Many TFs drive potent inflammatory responses that, when aberrantly activated, can contribute to inflammatory and autoimmune diseases. We therefore hypothesized that some TFs may require tight DNA-binding regulation. Indeed, transcriptionally activating and bifunctional TFs are typically subjected to more layers of specificity than repressing TFs or those with unknown transcriptional activity (**Fig. 5f**), suggesting that transcriptional activation is more tightly controlled than transcriptional repression. This is consistent with the requirement to avoid aberrant activation of cytokine expression, which can result in acute autoinflammation or chronic autoimmune disease. Furthermore, the five largest TF families represented in our pY1H network – NRs, bHLHs, C2H2 zinc fingers (ZFs), bZIPs, and homeodomains (HDs) – preferred distinct combinations of specificity mechanisms (**Fig. 5g**). Most NRs are controlled by all three mechanisms, consistent with their known expression specificity across immune cells, dependence on heterodimerization, and ligand binding. Most bHLH and HD TFs are associated with two layers of specificity. However, while bHLHs were typically activity- and partner-modulated, consistent with signal-dependent activity that is further mediated by TF partners, HDs were expression- and partner-modulated, suggesting a high level of cell type specificity that is consistent with the role of HDs in development and cell type differentiation. In both cases, partner modulation seems to complement previously reported mechanisms of specificity.

Given the detrimental effects of spurious or persistent cytokine activation, we hypothesized that TFs with major effects on cytokine expression and immune responses may be subjected to more layers of specificity. Indeed, TFs with more layers of specificity targeted a greater number of cytokine promoters in our pY1H screen (**Fig. 5h**). Importantly, the number of specificity layers is positively correlated with the proportion of TFs associated with mammalian immune phenotypes (**Figure 5I**). This suggests that many TFs which are essential for appropriate immune function are under tighter control by combinations of activity, expression, and partner modulation. Therefore, disruption of any of these mechanisms could potentially contribute to dysregulated cytokine expression and subsequent immunodeficiency or autoimmunity. For example, TNF*α* is a key proinflammatory cytokine whose aberrant expression is associated with autoimmune disease and cytokine storm^49,50^. We identified 19 TF-DNA interactions at the *TNF* promoter, which are subject to varying combinations of specificity mechanisms (**Fig. 5j**) (**Supplementary Table 14)**. Five of these PDIs – involving RUNX2, FOSL2, FOSL1, GLI3, and RARG – are limited by all three mechanisms; all five of these TFs have been associated with abnormal *TNF* expression^51-62^, and three (RUNX2, FOSL2, and FOSL1) are associated with immune-related phenotypes in the MGI database. This supports the idea that PDIs which can affect the expression of key pro-inflammatory cytokines are often controlled by multiple levels of regulation, mitigating the potential for aberrant expression by introducing AND gates to the interaction network.

## DISCUSSION

Our study provides a systematic view of how multiple regulatory layers - TF activity, expression, cooperativity, and antagonism - contribute to specificity in cytokine gene targeting. Although prior work has established that TFs can act combinatorially to drive cell type-specific programs, these conclusions have largely been drawn from indirect evidence of co-occupancy^22^ or studies of selected TFs^14,17,41^ or individual loci^63^. By integrating large-scale pY1H assay data with TF activation and expression data, we quantify the pervasiveness of these mechanisms and how they jointly constrain PDIs.

Our large-scale pY1H screens substantially expand the number of TFs and promoters connected within the cytokine network and recover a high fraction of PDIs supported by the literature or ChIP-based datasets. This complements prior studies of cytokine regulation by dimeric TFs such as NF-κB, STATs, and IRFs, which have been limited in their ability to systematically probe TF partner dependence. We found that many “promiscuous” TFs use different partners to engage distinct cytokine promoters, indicating that partner choice is a major determinant of DNA-binding specificity. Promiscuous TFs, including IKZF3, FOS, and MYC, are known hubs in immune differentiation and activation^17,31,64,65^. Our results suggest that the regulatory breadth of these TFs reflects not only their broad expression but also the availability of multiple TF partners which expand the repertoire of DNA targets and diversify the transcriptional effector domains recruited.

We further find that cooperativity frequently expands cofactor recruitment potential. Previous structural and biochemical work demonstrated that multi-TF complexes, such as those assembled in the interferon beta enhanceosome^63^, can increase affinity for coactivators and Mediator, amplifying transcriptional outputs. Similarly, cooperative pairs in our network have partially overlapping cofactor interaction profiles, supporting a model in which cooperative DNA binding both strengthens cofactor contacts and expands the repertoire of cofactors recruited. Many pairs also include one TF with no annotated effector domains or cofactor interactions, consistent with specialized relationships in which one TF tunes DNA binding while its partner mediates transcriptional output. Such relationships have been proposed for NF-Y TFs^66^ but have not been characterized at scale.

Surprisingly, we found that TF antagonism was also highly prevalent. Although antagonistic TF-TF relationships are well documented in specific systems^36^, such as inhibitory bHLH proteins^67^, the extent to which antagonism affects other TFs is unknown. Our pY1H assay data shows that antagonized interactions are nearly as common as those that rely on cooperativity. Many TFs engage in complex relationships in which a partner prevents binding to some targets while facilitating interactions with others, suggesting that TF-TF relationships exist on a continuum bridging pure antagonism, functional redirection, and pure cooperativity. This generalizes isolated mechanistic observations into a network-level view in which the effective binding of a TF to a promoter is strongly shaped by the local repertoire of its antagonists and cooperative binding partners. Interestingly, TFs that act purely as antagonists are just as likely to be linked to immune phenotypes as TFs that directly bind cytokine promoters, supporting the idea that disruptions in antagonistic partners can affect cytokine networks even when DNA-binding TFs are unaltered.

Our analysis of activity- and expression-modulated TFs places partner modulation (cooperativity and antagonism) in a broader context. Cytokine regulation is often attributed to inducible TFs whose activity is controlled by PTMs, ligand binding, or other mechanisms. While activity-modulated TFs are prevalent in both the literature-derived and Y1H-derived cytokine GRNs, Y1H data reveals that expression-modulated, cell type-specific TFs also frequently target cytokine promoters, which is consistent with the strong cell type specificity of cytokine expression. When all three layers—activity, expression, and partner modulation—are considered together, ∼97% of TF-promoter interactions appear to be constrained by at least one mechanism and ∼73% are constrained by two or three. This provides a mechanistic explanation for how restricted patterns of cytokine expression are achieved with often broadly expressed TFs: TFs often must be both expressed and activated, and simultaneously engage permissive partners while avoiding antagonistic ones. This multi-layered regulation of TF-DNA binding may provide added security in preventing spurious cytokine expression, which can cause acute and chronic autoinflammatory tissue damage. More targeted studies should explore how disruptions in these specificity mechanisms may contribute to cytokine dysregulation and autoimmune diseases.

Despite being restricted to the TF-pairs and promoters examined here, our findings indicate that TF partner modulation is likely a general mechanism shaping transcriptional regulatory specificity. Of the 328 cooperative binding events detected, 95 (29%) involved TF-pairs not previously known to co-bind DNA, suggesting that unbiased testing of additional TF-pairs is likely to reveal partner-dependent relationships. Furthermore, we previously used pY1H assays to reveal widespread cooperativity and antagonism at promoters of cancer-related genes^35^, demonstrating that these mechanisms are used to achieve specificity across gene families. However, TF-pairs were more than twice as likely to participate in cooperative and antagonistic events at cytokine promoters than they were at cancer gene promoters, consistent with the requirement for cytokine gene expression to be restricted to specific cell types and stimulatory conditions.

pY1H assays are limited to pairwise TF combinations and therefore cannot capture higher-order complexes involving three or more TFs, and because they are performed in yeast, they do not necessarily recapitulate native chromatin features such as nucleosome positioning, histone modifications, or DNA methylation that influence TF binding *in vivo*. However, this framework provides key advantages: by testing TF pairs in isolation, pY1H assays map pairwise interactions with minimal interference from other human TFs or cofactors, enabling the identification of interactions driven primarily by DNA sequence features and intrinsic TF properties. As such, these interactions define a baseline layer of sequence-encoded regulatory potential that can be further modulated by chromatin context in native cellular environments.

In summary, by interrogating single TFs and TF-pairs at cytokine promoters and integrating these data with activity and expression information, we show that TF specificity arises from a densely wired landscape of cooperative and antagonistic interactions layered on top of activity- and expression-based control. This framework helps explain how seemingly imprecise TFs achieve precise cytokine regulation and provides a resource for understanding how disruptions in TF networks contribute to immune dysfunction.

## METHODS

### pY1H screening

Interactions between TF-pairs and DNA-baits were screened as previously described^23^ and as follows. Using a high-density array ROTOR robot (Singer Instruments), the TF-pair yeast array and DNA-bait yeast strains were mated pairwise to test all possible combinations on agar plates with permissive media. Mated yeast were incubated at 30°C for one day and were then transferred to agar plates with selective media lacking uracil, leucine, and tryptophan, which only allow for growth of successfully mated diploid yeast. These selection plates were incubated at 30°C for two days, imaged, and analyzed to identify array locations with failed yeast growth, which were excluded from analysis. Diploid yeast were then transferred to agar plates with selective media lacking uracil, leucine, tryptophan, and histidine, with 5 mM 3AT and 320 mg/L X-gal. These readout plates were imaged after 2, 3, 4, and 7 days.

### Image processing

To analyze the pY1H images, we previously developed an open-source analyzer called DISHA (Detection of Interactions Software for High-throughput Analyses)^23^. Briefly, DISHA uses classical computer vision algorithms for boundary cropping and grid generation, followed by a UNet-based deep-learning model^68^ for colony segmentation. DISHA then computes colony area as the number of non-zero pixels within a segment, as well as the average intensity of blue coloration across the pixels within the colony. Finally, DISHA calculates a reporter signal score for each yeast strain, normalized to values from control yeast strains not expressing human TFs. These reporter scores are compared between yeast strains expressing TF1 only, TF2 only, or TF1+TF2 to calculate DNA-binding cooperativity and antagonism indexes^23^. DISHA also utilizes a visualization tool to allow users to view whole plates, easily compare relevant yeast strains, and view calculated reporter signal scores and cooperativity/antagonism indexes.

### Calling interactions

DISHA was used to sort TF-pair yeast strains by cooperativity index or antagonism index. Images were then assessed manually to call high-confidence cooperative or antagonistic events. To call a cooperative event, we required:

1. Successful growth of mated yeast on diploid selection plates for yeast expressing TF1 only, TF2 only, and TF1+TF2.
2. Uniform reporter signal for ≥3 out of 4 quadruplicate colonies for yeast strains expressing TF1 only, TF2 only, and TF1+TF2.
3. Either strong or moderate reporter signal for the yeast strain expressing TF1+TF2, and either weak, very weak, or no signal for strains expressing TF1 only and TF2 only.

To call an antagonistic event in which TF1 antagonized TF2, we required:

1. Successful growth of mated yeast on diploid selection plates for yeast expressing TF2 only and TF1+TF2.
2. Uniform reporter signal for ≥3 out of 4 quadruplicate colonies for yeast strains expressing TF2 only and TF1+TF2.
3. Either strong or moderate reporter signal for the yeast strain expressing TF2 only, and either weak, very weak, or no signal for the strain expressing TF1+TF2.

Called interactions can be found in Supplementary Tables 4, 5, 8, and 9.

### Overlap between ChIP-seq and pY1H interactions

To identify evidence for pY1H-derived TF-promoter interactions in human cells, we obtained ChIP-seq data from the GTRD database^69^ (peaks calling = MACS2, reference genome = hg38, format file = bigBeds). We identified significant peaks (p-value ≤ 10^-4^) for which the summit was located anywhere within the ∼2kb promoter sequence tested by pY1H. We considered the TF-promoter interaction to have ChIP-seq evidence if ≥1 such peak was identified in ≥1 cell line.

To determine whether pY1H and ChIP-seq interactions overlap more than expected by chance, we generated 100,000 randomized networks using the nodes from the pY1H network and performed 20,000 edge-switches. We then generated a distribution of the degree of overlap between these 100,000 randomized networks and ChIP-seq data. Finally, we compared this distribution to the degree of overlap between the pY1H network and ChIP-seq data to obtain a Z-score and two-tailed p-value.

Code for ChIP-seq analysis can be found here: https://github.com/jfuxman/PY1H_NatComm2023/tree/main/Obtaining%20ChIP-seq%20data%20from%20GTRD

Code for randomization analyses can be found here: https://github.com/jfuxman/PY1H_NatComm2023/tree/main/Network%20Randomization%20Analysis%20(EY1H%20and%20PY1H%20with%20ChIP%20data)

The number of pieces of ChIP-seq evidence for each pY1H-derived PDI can be found in Supplementary Table 4.

### Data visualization and statistical analyses

Network schematics were generated using Cytoscape Version 3.10.3. Scatter plots, violin plots, histograms, bar graphs, and heat maps were generated using GraphPad Prism Version 10.

### Identification of known and novel co-binding TF-pairs

TF-pairs with at least one cooperative binding event observed by pY1H assays were assessed to determine whether the two TFs had been previously shown to co-bind DNA. Two graduate-level scientists searched the literature using PubMed, Google Scholar, and Google Images for reports that mentioned both of the TFs and DNA binding. A TF-pair was considered to be a known co-binder of DNA if at least one piece of evidence was found for simultaneous or cooperative DNA binding, binding as a heterodimer, or co-occupancy in DNA-binding complexes.

Annotations of known and novel co-binding pairs can be found in Supplementary Table 5.

### Overlap between cofactor interaction profiles

We compiled a list of known interactions between TFs in cooperative TF-pairs and transcriptional cofactors using the BioGRID database^32^ downloaded on August 13, 2025. A TF-cofactor interaction was considered if it had ≥3 pieces of evidence reported in BioGRID. For each cooperative TF-pair, we calculated the Simpson similarity index of their cofactor interaction profiles. We then generated a list of 3900 randomized TF-pairs using the TFs included in the list of cooperative TF-pairs. We compared the distributions of Simpson similarities for cooperative and randomized TF-pairs using a two-tailed Mann-Whitney U test.

TF-cofactor interactions can be found in Supplementary Table 7.

### TF and cytokine expression analysis

Expression data for TFs and cytokine genes was obtained from the Tabula Sapiens single cell RNA-seq atlas^28^ and analyzed as previously described^23^. Briefly, we compiled data obtained using 10X Genomics protocols from cells with 500-7500 genes expressed, ≤ 10,000 unique molecular identifiers detected, and ≤ 25% of reads originating from mitochondrial genes. We converted read counts to log-normalized counts per million (cpm), performed principal component analysis, and mitigated batch effects using Harmony^70^. We used a k-nearest neighbor graph and Louvain community clustering to identify 187 cell clusters, each of which was assigned a known cell type by comparing enriched genes to cell type markers annotated in CellTypist^71^.

For expression profile similarity and overlap analysis, we considered a gene to be “expressed” in a given cell type if the cpm value for the gene in that cell type was higher than 10% of the maximum cpm value for the gene across all 187 cell types.

Code for assessing TF and cytokine gene expression from the Tabula Sapiens atlas can be found here: https://github.com/jfuxman/PY1H_NatComm2023/tree/main/TF%20expression%20analysis

Expression values can be found in Supplementary Table 6.

### Tissue/cell type expression specificity scoring of genes

Using expression values across cell types identified using data from Tabula Sapiens, we calculated a tissue/cell type expression specificity score (TCESS) for each TF and cytokine gene in the cytokine GRN as previously described^9,23^. Briefly, for an example gene TF_a_, given a cluster C containing n cells, the total expression of TF_a_ was calculated using the following formula:

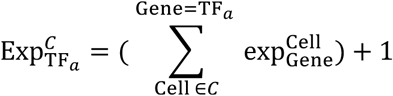

The TCESS was then calculated as follows:

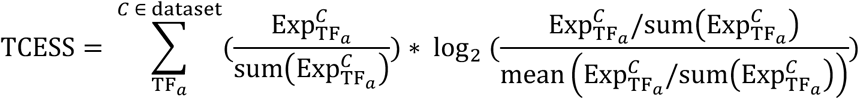

The TCESS ranges from 0 (where gene expression is identical across all cell types) to log_2_(#cell types), in this case ∼7.54, (where a gene is expressed exclusively in one cell type).

Code for calculating TCESS scores can be found here: https://github.com/jfuxman/PY1H_NatComm2023/tree/main/TF%20expression%20analysis

TCESS values can be found in Supplementary Table 12.

### Annotation of TFs as potential activators, repressors, bifunctional, or unknown

We compiled a list of known TF effector domains (activation, repression, and bifunctional domains) by combining annotations from Soto *et al.*^33^ and high-throughput reporter assay results from DelRosso *et al.*^34^. TFs containing ≥1 activation domain, 0 repression domains, and 0 bifunctional domains were considered activators. TFs containing 0 activation domains, ≥1 repression domain, and 0 bifunctional domains were considered repressors. TFs containing ≥1 activation domain and ≥1 repression domain or containing ≥1 bifunctional domain were considered bifunctional. TFs with no known effector domains were considered unknown.

Annotations for TFs can be found in Supplementary Table 2.

### Mammalian immune phenotypes and human immune diseases

We used the Mouse Genome Informatics (MGI) database to identify associations between TFs and 1190 mammalian phenotypes in the “immune system phenotype” category curated by MGI. We also used MGI to search for associations between TFs and human diseases, manually curating a subset of 21 human diseases associated with immune function. We considered a TF to be associated with mammalian immune phenotypes or immune-related human diseases if at least one piece of evidence was provided in MGI.

Associations between TFs and mammalian immune phenotypes or human immune diseases can be found in Supplementary Tables 10 and 11.

### Classification of activity-modulated TFs

We determined whether each TF in the cytokine GRN could be modulated by dynamic changes in activity. We considered a TF to be activity-modulated if it met at least one of the following criteria:

1. Binding of a ligand is known to modulate the transcriptional activity, DNA binding, or nuclear localization of the TF.
2. A post-translational modification (phosphorylation, methylation, acetylation, or sumoylation) is known to modulate the transcriptional activity, DNA binding, or nuclear localization of the TF. This information was mainly derived from annotations in PhosphoSitePlus^10^ and supplemented with searches in PubMed and Google Images.
3. Localization of the TF to the nucleus or to DNA is otherwise dynamically regulated, e.g. by sequestration of the TF in the cytoplasm or at the nuclear lamina or by proteolytic cleavage of the TF.

Annotations for TFs can be found in Supplementary Table 12.

### Classification of expression-modulated TFs

We considered a TF to be expression-modulated if it met at least one of the following:

1. The TF was detected but not ubiquitously expressed in the Tabula Sapiens single-cell (sc)RNA-seq atlas^28^. We required a tissue/cell type expression specificity score (TCESS) of >0.25, similar to the threshold between ubiquitous and cell type-specific TFs selected by Ravasi *et al.*^9^.
2. The TF showed changes in expression in response to extracellular signals in the CytoSig database^48^. For a TF to be considered expression-modulated in response to signals, we required that its RNA expression was either upregulated (log_2_(fold change)>2) or downregulated (log_2_(fold change)<-2) in more than 20 (∼1%) experiments with data in CytoSig.

Annotations for TFs can be found in Supplementary Table 12.

### Association between TFs and TNF expression

To identify reported associations between TFs and TNF expression, we searched the literature using PubMed, Google Scholar, and Google Images to find papers mentioning the TF and TNF. We manually curated relevant papers to identify those with direct evidence connecting perturbation of the TF with changes in TNF expression at the RNA or protein level. We also incorporated associations reported in the MGI database.

Information can be found in Supplementary Table 14.

## Supporting information

Combined Supplementary Tables

## ACKNOWLEDGEMENTS

This work was funded by the National Institutes of Health grant R35 GM128625 awarded to J.I.F.B.

## AUTHOR CONTRIBUTIONS

A.L. and J.I.F.B. conceived the project. A.L., R.L., and S.S. performed the pY1H screens. A.L., Y.L., Z.L., L.S.-U., and M.A.P. performed data analyses. M.P. and C.C. developed DISHA. A.L. and J.I.F.B. wrote the manuscript. All authors read and approved the manuscript.

