## Supplementary figures and images for "Multilayered specificity of transcription factor binding at cytokine promoters"

### Supplementary Figure 1

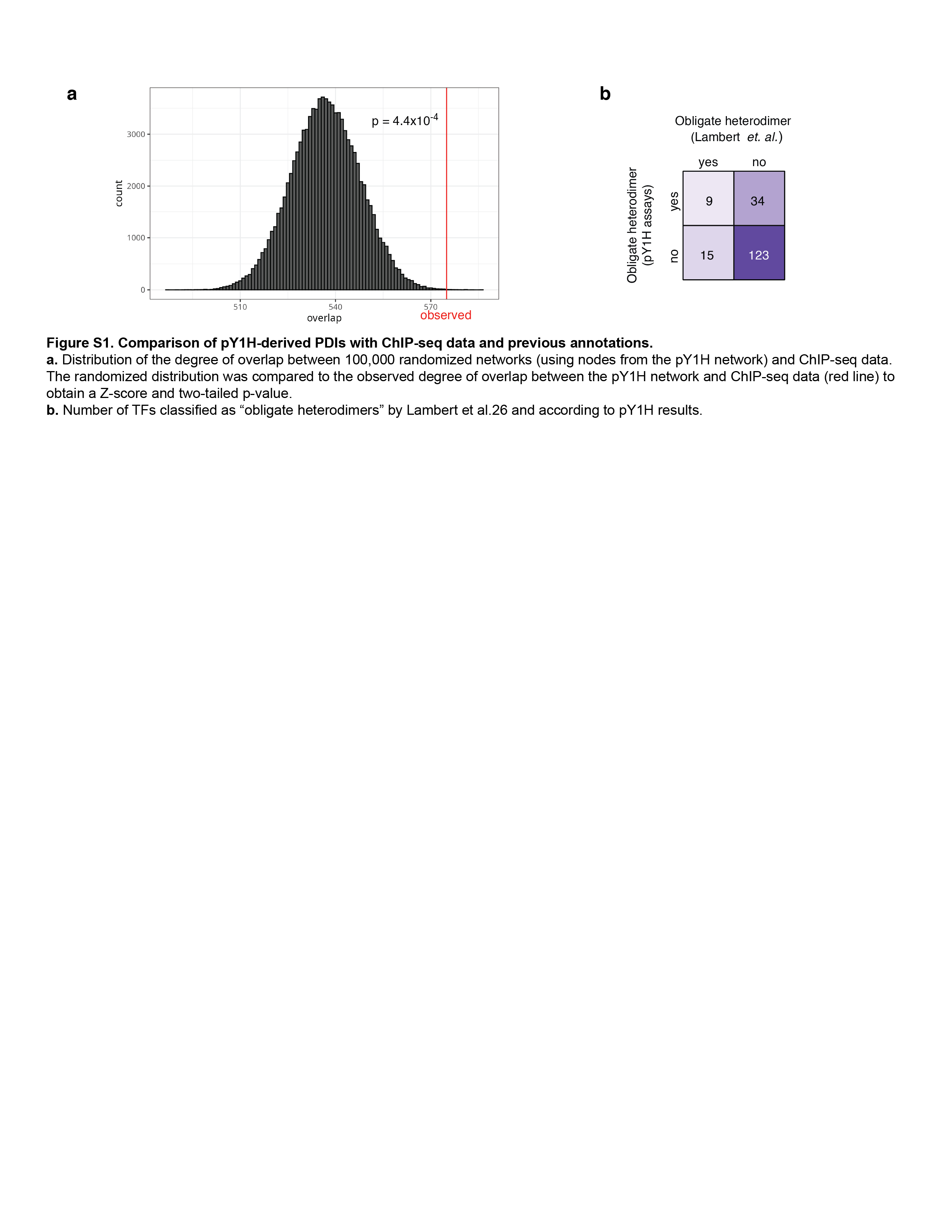

### Supplementary Figure 2

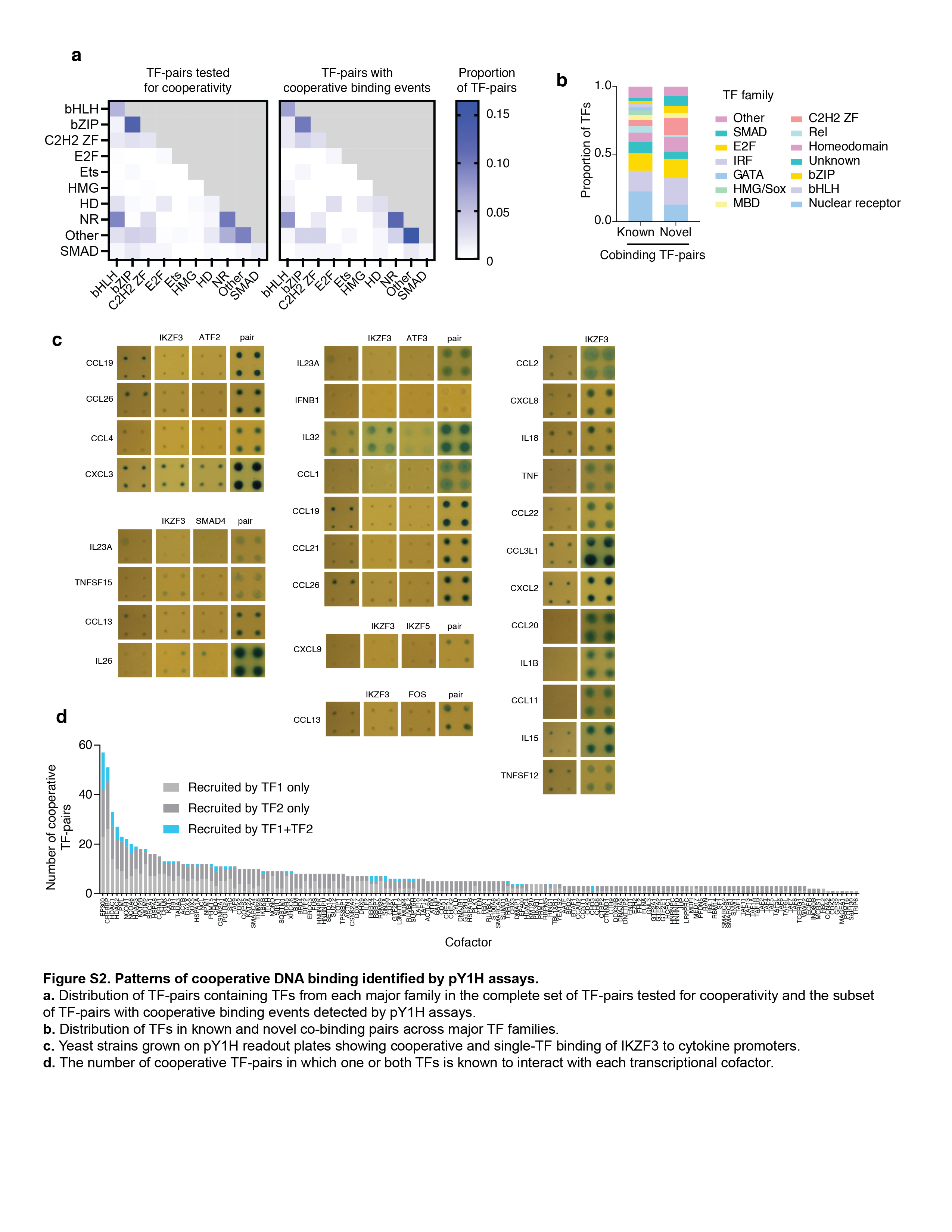

### Supplementary Figure 3

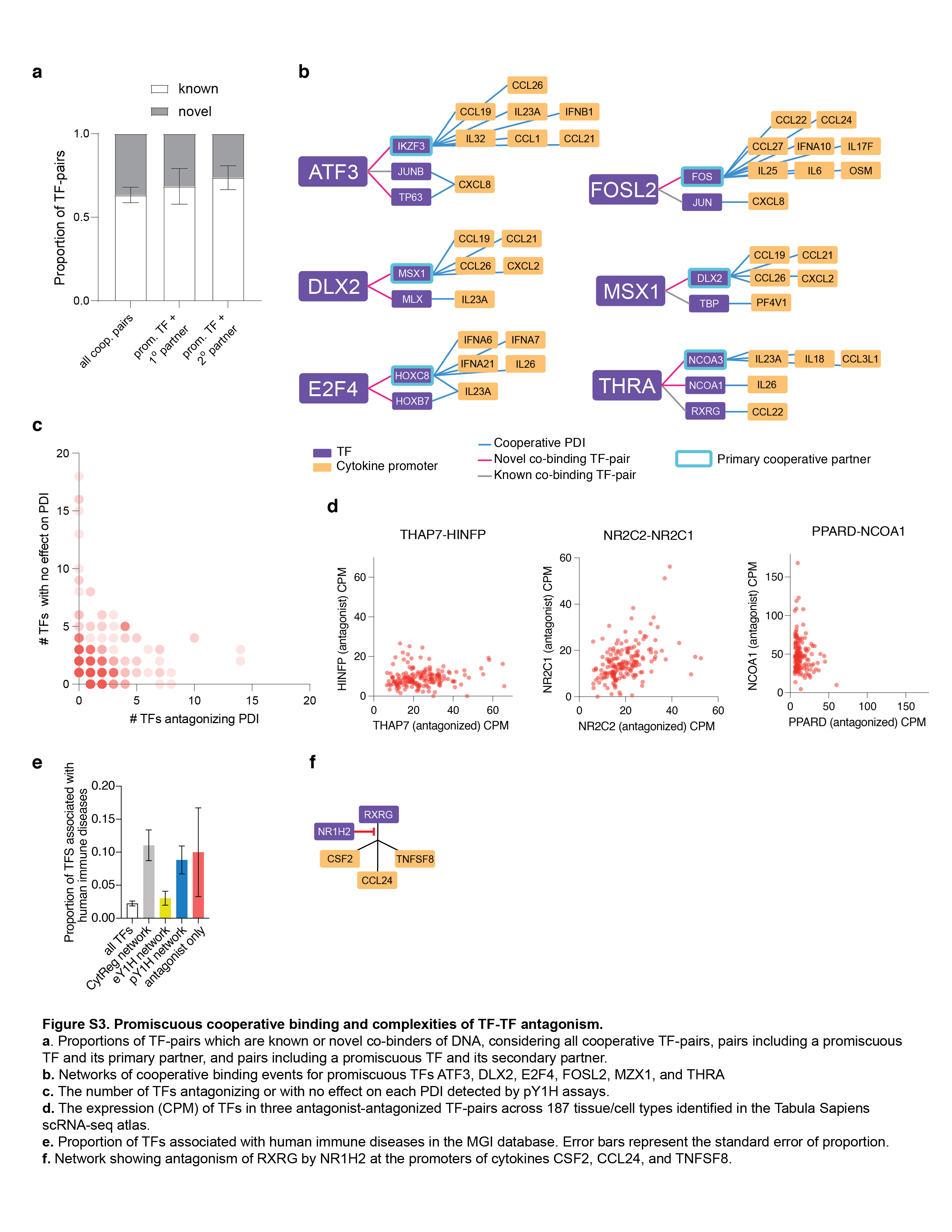
